# Multi-frequency tACS has lasting effects on neural synchrony and audiovisual responses

**DOI:** 10.1101/526087

**Authors:** Tobias Bockhorst, Joachim Ahlbeck, Florian Pieper, Gerhard Engler, Andreas K. Engel

## Abstract

**BACKGROUND:** In cortical networks, synchronized oscillatory activity of neuronal populations enables communication, and its disturbance is related to a range of pathologies. Transcranial alternating current stimulation (tACS) has been explored as a flexible, noninvasive tool for the modulation and restoration of synchronized oscillatory signals. While numerous studies have addressed cognitive or behavioural effects of electrical stimulation, the neural changes underlying the effects of tACS and their persistence after stimulation offset have remained unclear.

**OBJECTIVE:** Here, we screened for lasting aftereffects of prolonged tACS on intrinsic network activity and audiovisual processing in the anesthetized ferret brain.

**METHODS:** Electrical stimulation was applied via subcutaneous wire electrodes. Current waveforms were synthesized from two frequencies in the alpha or gamma range, respectively. Flashes and clicks were used for audiovisual stimulation. Electrocorticographic recordings from an extended network including occipital, temporal and parietal cortical areas were obtained before and after tACS.

**RESULTS:** Changes in local synchrony (continuous and spike-triggered power of LFP), synchrony across recording sites (imaginary coherence) and altered dynamics of sensory response-features (peak-to-peak amplitude, extremum latency) following electrical stimulation consistently point to a synchronizing effect of tACS that can outlast stimulation offset by at least 10 min. Gamma-band tACS proved particularly effective. In line with previous reports on cross-frequency interactions, we observed effects on coherence and power of baseline activity at frequencies other than the ones targeted by tACS. These cross-frequency interactions appeared to underlie the strengthening and stabilizing effect on audiovisual responses.

**CONCLUSION:** We demonstrate aftereffects of tACS on synchrony and stimulus processing in an extended cortical network, measured intracranially in a setting that resembles tACS stimulation in humans. The data provide direct evidence for the efficacy of tACS as a tool for sustained modulation of cortical network dynamics.

## Introduction

Oscillatory brain signals reflect the coordinated activity of large neuronal populations [1]. Synchrony of oscillations within and between brain regions is considered a fingerprint of functional coupling underlying perceptual, sensorimotor and cognitive processing [2–8]. Impairment of proper coupling may lead to disorders, such as AD, PD, epilepsy, schizophrenia and autism [9]. This has fueled research into non-invasive approaches to modulate and, possibly, re-adjust neural synchrony by transcranial alternating current stimulation (tACS) [10–14].

Most studies have focused on direct effects during electrical stimulation, and key advances include focal stimulation of deep brain targets [15, 16], the induction of artificial percepts that drive behavior [17], suppression of tremor in Parkinson’s disease [18] and automated dampening of epileptic seizures [19]. Modulation of percepts or detection thresholds during tACS have also been reported in several studies [20–23].

In contrast, relatively little is known regarding persistent physiological effects of tACS, i.e., aftereffects on neural activity that outlast the period of electrical stimulation. Lasting modulation of intrinsic oscillations (predominantly in the alpha-band) and persistent effects on task performance or mental state have been reported in noninvasive studies [14, 24–27]. By contrast, studies addressing electrophysiological aftereffects of tACS by invasive recordings have not yet been performed. Furthermore, electrophysiological evidence from in-vivo studies employing stimulation with continuous waveforms has been restricted to local activity captured at single recording sites [17, 28]. As a result, effects of tACS on the state of cortical networks, key to clinical application in disorders of brain connectivity, remained largely unaddressed.

Here, we analyzed unilateral electrocorticographic recordings from an extended network including occipital, temporal and parietal cortical areas in the ferret, to screen for persistent effects of tACS on intrinsic and stimulus-related activity. In addition to stimulation in the most commonly investigated alpha range, we included tACS at higher frequencies in the upper beta and gamma band. To reasonably match the situation in humans, tACS was applied subcutaneously, leaving intact the particularly voluminous muscles of mastication as well as the skull. In anatomical respects, the gyrification of the ferret cortex (unlike rats and mice) parallels the situation in humans, which factors-in effects of anisotropy on current flow and relative orientation of electric fields to somato-dendritic axes of the neurons located in gyri and sulci.

## Material and Methods

### Subjects

Four adult female ferrets (*Mustela putorius furo*) were used for experiments (Euroferret, Dybbølsgade, Denmark). Ferrets were kept in standardized ferret cages (Tecniplast, Hohenpeißberg, Germany) with an enriched environment under controlled ambient conditions (21°, 12-h light/dark cycle, lights on at 8:00 a.m.). The animals had ad libitum access to food pellets and tap water. All experiments were approved by the independent Hamburg state authority for animal welfare (BUG Hamburg) and were performed in accordance with the guidelines of the German Animal Protection Law.

### Surgery

Details on the surgical procedure have been reported earlier [29]. Briefly, animals were initially anesthetized with an injection of ketamine (15 mg/kg) and medetomidine (0.08 mg/kg). A glass tube was inserted into the trachea to allow artificial ventilation of the animal and to supply isoflurane (approx. 0.7-0.84 %; exact concentration adjusted as to avoid burst-suppression states) for maintenance of anesthesia. N_2_O (30%) was added to the ventilated air for its analgesic and anesthetic effects. Body temperature was monitored rectally and automatically maintained at 37.5°C. Heart rate and end-tidal CO_2_ concentration were constantly monitored throughout the whole experiment to maintain the state of the animal. To prevent dehydration, a cannula was inserted into the femoral vein to deliver a continuous infusion of 0.9 % NaCl and pancuronium bromide (6 μg/kg/h). Contact lenses were placed on both eyes to avoid desiccation of the cornea.

The temporalis muscle was gently detached and pushed aside, such that a craniotomy could be performed over the left posterior cortex using a saline-cooled and soft-tissue-safe micro-saw. The dura was incised and folded back to allow placing the ECoG array on the surface of the cortex. Once the array’s positioning had been adjusted as to best cover all cortical areas of interest, the dura was relocated back and supplemented with artificial dura (Lyoplant; B. Braun, Melsungen, Germany) as to cover the entire ECoG array. Finally, the craniotomy was closed by re-implanting the previously removed piece of parietal bone plate, which was kept in saline before to prevent an increase in ohmic resistance, sealing the rim with KWIK-CAST silicon-based glue (World Precision Instruments), folding back the temporalis muscle into anatomical position and sewing up the skin incision.

### Custom ECoG design

Recordings were carried out using a custom-made ECoG array that matches the anatomy of the left hemisphere of the ferret brain. The array consists of 64 platinum electrodes with a diameter of 250 μm each, embedded in a thin-film polyimide foil of 10 μm thickness (7-25 kΩ @ 1kHz) and arranged in a hexagonal formation with an interelectrode distance of 1.5 mm [29,30].

### Recording

All experiments were performed on a vibration-damped table (Technical Manufacturing Cooperation, Peabody, MA, USA) inside a soundproof and electromagnetically shielded booth (Acoustair, Moerkapelle, Netherlands). Neural signals were band-pass filtered between 0.5 Hz to 8 kHz, digitized at 44 kHz / 16 bits and sampled with an AlphaLab SnR™ recording system (Alpha Omega Engineering, Nazareth, Israel). All analyses of neural data presented in this study were performed offline after the completion of experiments using custom-written Matlab scripts (Mathworks). For spike detection, both ECoG and intracortical signals were band-pass filtered between 300 Hz and 5 kHz, transformed to zero median and normalized to the long-tail-corrected standard deviation (median of the absolute values divided by 0.6745; according to [31] of the time series. Spike times were identified as local minima below a threshold of −3 standard deviations (artifact rejection threshold: −20 standard deviations), with no refractoriness assumed for the multi-unit activity measured here.

### Sensory stimulation

Sensory stimuli consisted of full-field flashes (100 ms duration, LED bright white light) framed by acoustic clicks (100 μs, broad-band) at flash ON and flash OFF. Stimulus blocks varied in length between experiments, covering about 400 - 600 trials (inter-stimulus-intervals 1-2.5 sec).

### Electrical stimulation

For tACS, wire electrodes (silver-wire, Science Products, Hofheim, Germany, 500 μm diameter, ~10 mm length, chlorinated; 1-3 kΩ impedance at 1 kHz) were implanted subcutaneously; positioned anterior and posterior to the area covered by the ECoG array. Alternating-current waveforms were generated by linear summation of two sinusoids with individual frequencies of 8 and 11 or 25 and 36 or 55 and 80 Hz; selected to modulate power in the alpha, higher beta or gamma band, respectively. Stimulation was applied for 40 minutes at 1 mA current (peak to peak amplitude) generated by a battery-driven, remote controlled stimulator (DC-Stimulator PLUS, neuroConn, Ilmenau, Germany). To exclude any electrical interference from the device, the stimulator was unplugged from the stimulation electrodes and removed from the shielded booth right after tACS.

### Analysis

For each animal, the position of ECoG arrays over posterior cortex was documented during surgery by taking photographs through a Zeiss OPMI pico microscope. The position of all 64 ECoG electrodes was then projected onto a scaled model ferret brain. The precise position of a reference electrode on the scaled ferret brain was measured, along with the angle required to rotate the ECoG such that all electrodes aligned with the photography taken during surgery. A rotation matrix was then used to translate the location of all 64 ECoG electrodes onto the standard map of the ferret cortex [29,32,33]. The cortical region underlying each electrode was then assigned to a cortical area and further assigned to one of 5 higher order groups as follows: EVis (Area 17, 18, 19), HVis (Area 20, 21, SSY), PP (PPC, PPr), SoSe (SII, SIII, 3b) or Aud (PSF, PPF, A1, AAF, AVF, ADF, fAES).

Power spectra were calculated using continous wavelet transform; coherence spectra were calculated using the imaginary part of the coherence spectrum [34] to exclude zero time-lag synchronization due to volume conduction. To assess tACS aftereffects on response strength, amplitudes of response waveforms (3-200 Hz; peak-to-peak) were first categorized as high-state (low-state), when being above (below) the half-maximum amplitude in the respective stimulus block, computed after convolution of the amplitude time-series with 25 elements boxcar-kernel. Non-parametric permutation based statistics were applied (10000 repetitions) to identify affected frequency ranges for power and coherence as well as the significance of response--amplitude state durations. For illustration of the permutation-based significance test procedure, see (Fig. S4). Non-parametric rank-sum tests were applied for pairwise comparisons of distributions (Bonferroni-corrected).

## Results

### Baseline sensory responses in the cortical network

We recorded local field potentials (LFP; 1 – 300Hz) and spiking activity from the cortical surface in four anesthetized ferrets that were implanted with a 64 channel electrocorticographic (ECoG) array [29,30,35]. The array was positioned to cover sensory and associative areas, including early visual cortex (EVis), higher visual cortex (HVis), posterior parietal cortex (PP), somatosensory cortex (SoSe), and auditory cortex (Aud) (see Methods for details) [29,32,33] (Fig.1A,B).

**Fig. 1.**
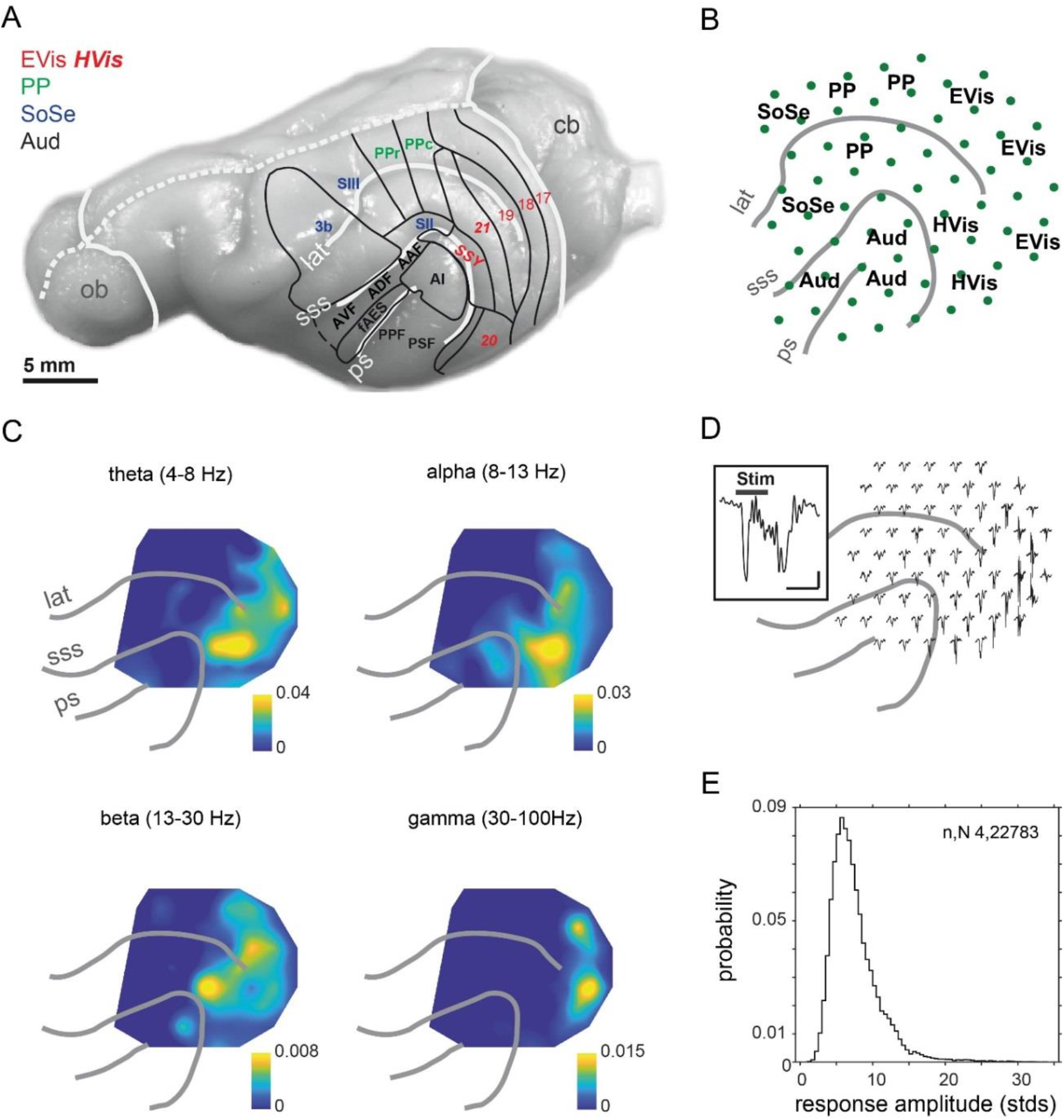
Electrocorticographic measurement of responses to audiovisual stimuli. **(A)** Brain areas covered by epicortical recordings in the resent study. EVis, early visual (areas 17-19); HVis, higher visual (areas 20 & 21; SSY, suprasylvian field); PP, posterior parietal (PPr, rostral posterior parietal cortex, PPc, caudal posterior parietal cortex); SoSe, somatosensory (area 3b; SII, SIII); Aud, auditory (AI; ADF, anterior dorsal field; AVF, anterior ventral field; AAF, anterior auditory field; fAES, anterior ectosylvian sulcal field; PPF, posterior pseudosylvian field; PSF, posterior suprasylvian field). ob, olfactory bulb; cd, cerebellum; white lines, prominent sulci (lat, lateral sulcus; sss, suprasylvian sulcus; ps, pseudosylvian sulcus; dashed line: midline). **(B)** Projection of recording electrodes onto the cortical surface, with labels of brain regions as denoted in (A). **(C)** Individual subject’s example topographies of increases in band-specific power during flash-click stimulation. (500ms windows located right before and after onset of stimulus; first 100 trials in block). **(D)** Example topography of trial-averaged response-waveforms; inset: detailed single-trial example; stim: stimulus; bars: 100ms, 1 μV. (same individual as in (C)) **(E)** Peak-to-peak amplitudes (in unit standard deviation) of single-trial responses (4 subjects, 22783 responses, first-of-day measurements only).

Each ferret was presented with several blocks of audiovisual “flash-click” stimuli (400-600 stimuli, see Methods). These evoked responses at sensory and parietal sites, accompanied by increased LFP power in theta, alpha and beta ranges (Evis, Hvis, PP, Aud), as well as in the gamma band (solely EVis) (Fig. 1 C-E). Somatosensory areas showed weakest responses (comp. Fig. 1B,D). Strongest responses to the audiovisual stimulus were recorded from visual areas (EVis, HVis) (Fig. 1D), with relative response amplitudes of up to 20 standard deviations (Fig. 1E). Taken together, the flash-lick stimulus proved adequate to study possible effects of tACS on sensory processing at the local and network level.

### Aftereffects of tACS on neural synchrony are frequency-specific

Current of 1mA (peak to peak) was applied for 40 minutes through subcutaneous wire electrodes positioned anterior and posterior to the area covered by the ECoG array (see Methods), with current waveforms composed of two superimposed sinusoids in the alpha band, higher beta band (one experiment) or gamma band, respectively (8 & 11 Hz, 25 & 36 Hz or 55 & 80 Hz) (Fig. 2A-C). Lower stimulation currents (500mA, 750 mA) were occasionally tested as well but did not prove effective. Audiovisual stimuli were presented prior and subsequent to tACS (Fig. 2D).

**Fig. 2.**
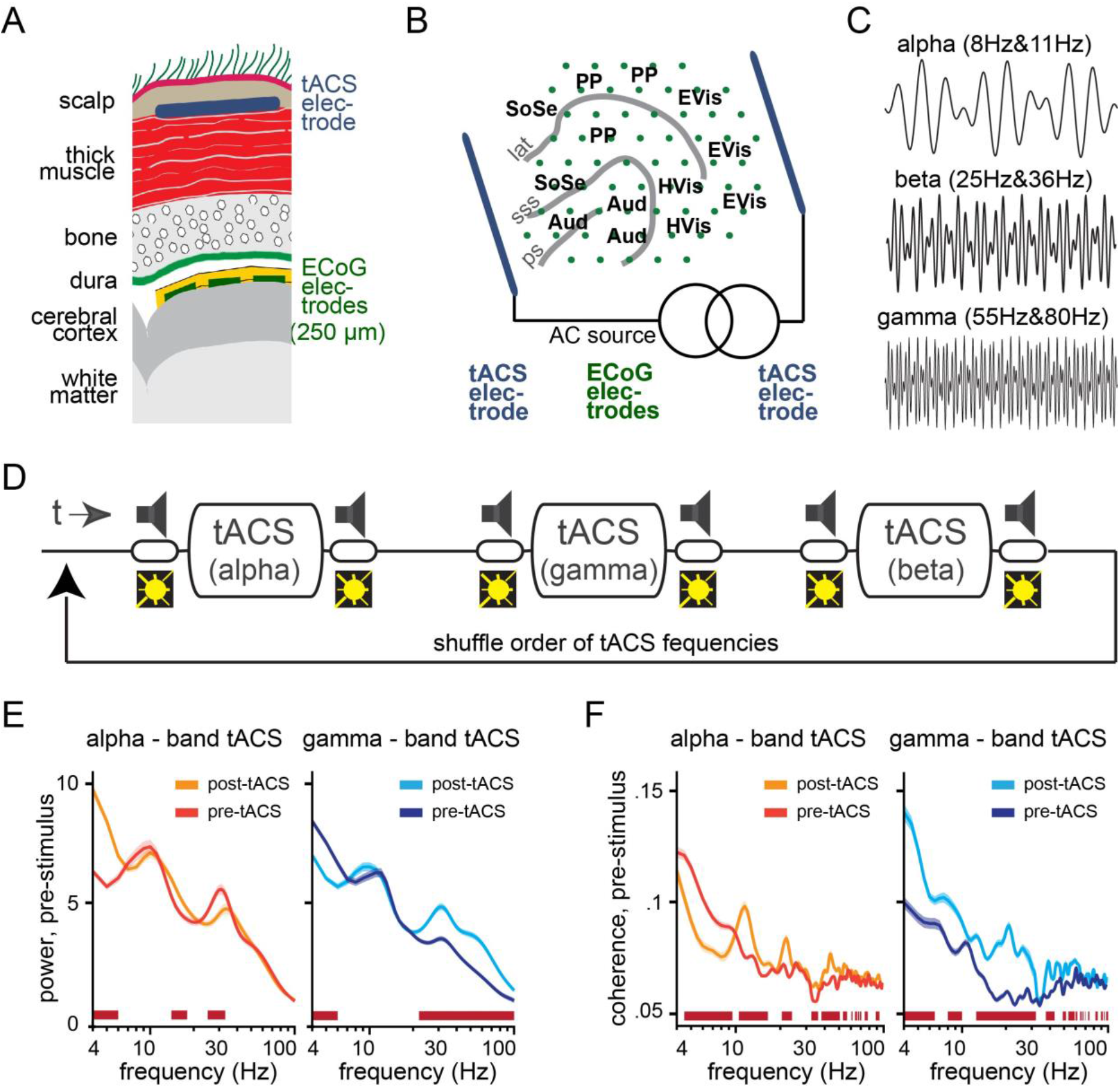
Effects of transcranial current stimulation: intrinsic activity. **(A)** Laminar placement of subcutaneous stimulation (tACS) and subdural recording (ECoG) electrodes. **(B)** Projection of recording and stimulation electrodes onto the cortical surface. Abbreviations as in Fig.1a. **(C)** Stimulus waveforms of alpha-, beta- and gamma-frequencies tACS (one second segments, synthesized by linear summation of the two respective frequencies specified above). **(D)** Schematic of experimental sessions. In each session, the order of tACS frequencies was shuffled as to vary both across animals and, within the same animal, between repetitions of the illustrated “triplet block”. Presentations of flash-click stimuli (400-600 trials per block, inter-trial-interval 1-1.5 sec) occurred both prior to and directly after tACS. Note that the audiovisual stimulus block prior to the very first application of tACS (first-of-day control) is not indicated. **(E)** Power spectra measured prior (pre) to and after (post) application of alpha- (/gamma-) band tACS (500 ms pre-stimulus window; first 100 trials in block; (all 64 recording sites considered; alpha: 3 experiments in 2 subjects;gamma: 5 experiments in 4 ferrets). **(F)** Coherence spectra, same general logic as in (E). Red horizontal lines in (E), (F) indicate statistical significance (10000 permutations).

Application of tACS caused changes in power and coherence spectra obtained from activity in baseline epochs (500 ms) preceding the sensory stimuli. More precisely, alpha-band tACS increased power in the theta range (4-6 Hz), while reducing power at frequencies in the beta range. A significant effect on power in the targeted alpha-band itself was not observed. Gamma-band tACS had a different effect, strongly increasing power in the targeted gamma-range and up to 100Hz (Fig. 2E), while reducing low-frequency power. Thus, gamma-band tACS lead to a shift in the ratio between gamma and theta power. Furthermore, both tACS regimes had different effects on coherence, i.e., the degree to which oscillations were synchronized across recording sites. Here, alpha-band tACS decoupled activity below 10Hz, while increasing coherence at some higher frequencies (around 12, 21 and 45 Hz). In contrast, gamma-band tACS broadly increased coherence across frequencies up to the high gamma range (80Hz), whereby the relative increase was larger in the gamma-than in the alpha-range (Fig. 2F).

### Coherence-modulation coincides with stabilizing aftereffects on audiovisual responses

We next asked whether tACS had also produced a lasting effect on the processing of the audiovisual stimuli. Here, a subset of recording sites, which clustered anatomically (**response-modulated sites**; Fig. 3A,B; Fig.S1), showed significantly higher power of stimulus-induced activity (up to 40Hz) as well as stronger cross-site coupling (up to 30Hz) (Fig. 3C,D). For these response-modulated sites, the time series of response amplitudes and latencies were strikingly different from those at recording sites which had not been affected by tACS (**non-modulated sites**) (Fig. 3E-I). This effect was also more readily observed after gamma-band than alpha-band tACS (Table 1, Fig. S1), possibly as a result of the abovementioned broader and stronger increase in neural synchrony.

**Fig. 3.**
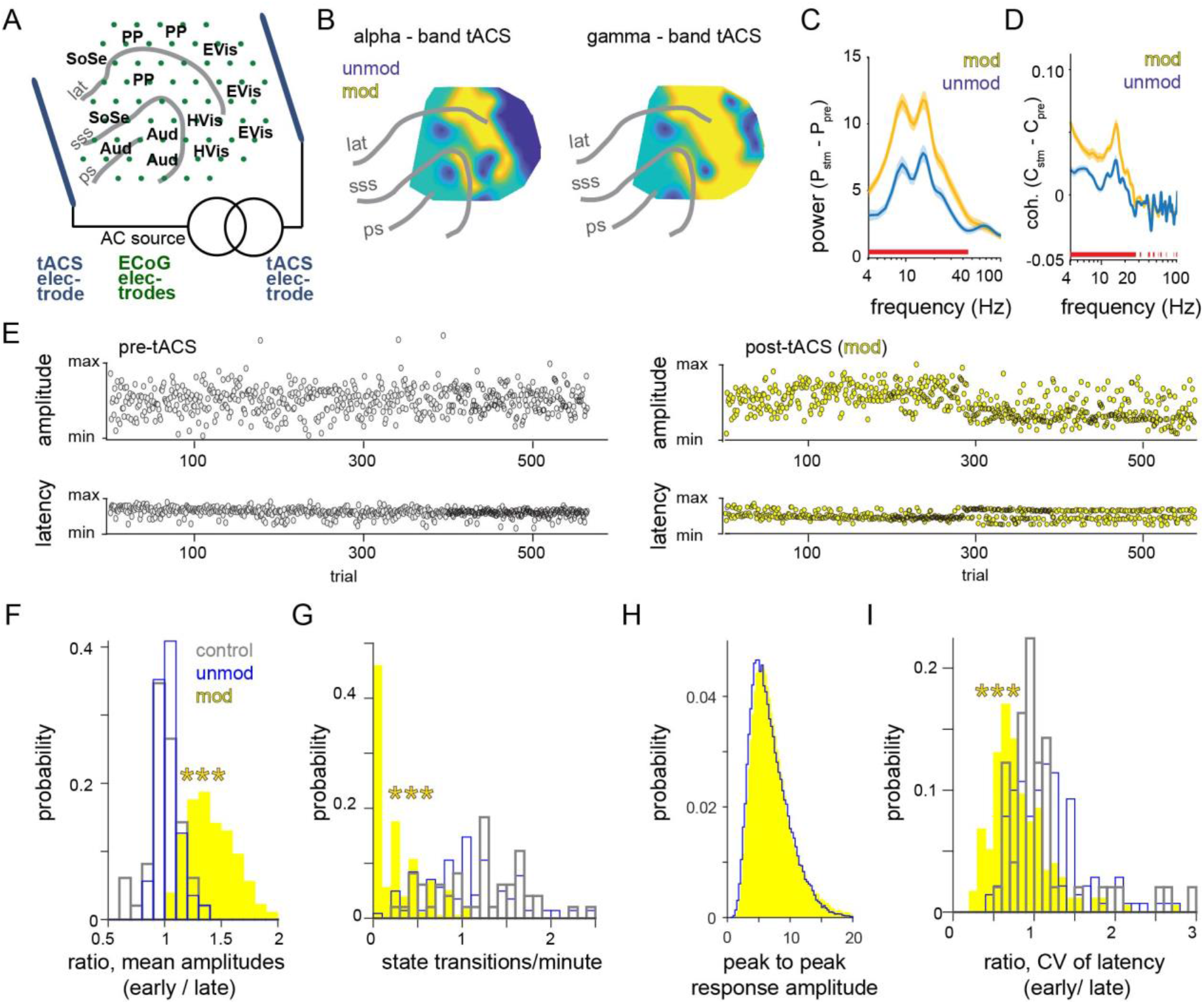
Effects of transcranial current stimulation: audiovisual responses. **(A)** Anatomical mapping of the measurement and stimulation electrodes. Abbreviations as in Fig.1a. **(B)** Example topographies of response-modulated vs. unmodulated sites after alpha- and gamma-band tACS. **(C,D)** Stimulus-related increase in mean power and imaginary coherence (500 ms windows; post-stimulus minus pre-stimulus; first 100 trials in block; averaged across tACS bands). Red horizontal bar indicates significance in difference between modulated vs. unmodulated sites (10.000 permutations). Shadings indicate spread (SEM). **(E)** Example time-courses of response amplitude and latency for a pre-tACS control (left) vs. post-tACS measurement at a modulated recording site (inter-trial interval 1-2.5s). **(F-I)** Distributions of mean amplitude ratio, frequency of transitions between high-amplitude and low-amplitude state, peak to peak response amplitudes and variance-of-latency ratio. CV: coefficient of variation; early, late: first, last 100 trials. Number of recording sites included: pre-tACS control: 49; unmod: 142; mod: 176.

**Table 1.**
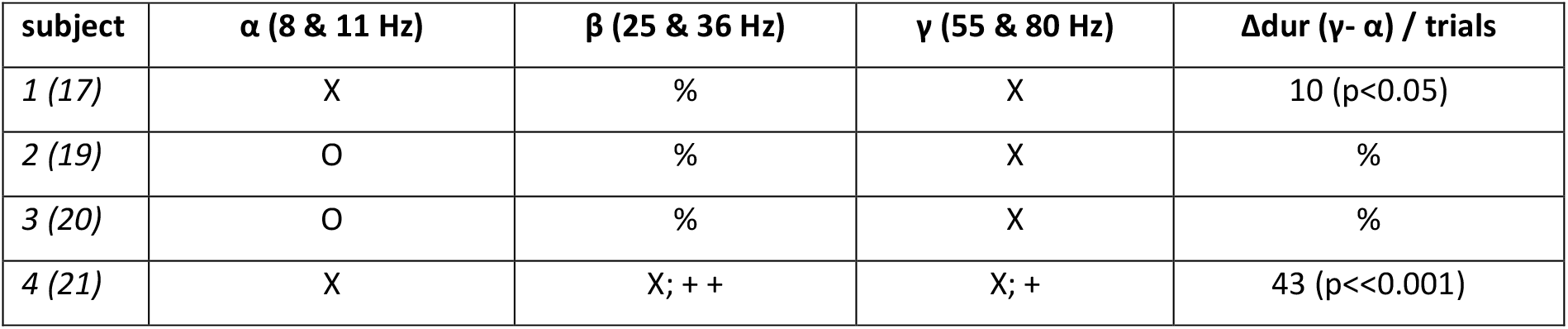
Effectiveness of different tACS frequency-pairs across experiments; O:= tested, ineffective; x: = tested, effective; %= not tested / not available; ++: = longest effect.; Δdur: difference (in trials) in duration of initial high state effect after tACS.

Prior to tACS, as well as at non-modulated sites after tACS, response amplitudes fluctuated across trials in a rather rapid manner (Fig. 3E), with transient states of high- and low-amplitudes alternating up to twice per minute (Fig. 3 F,G). Maximum state durations amounted to 2.5 – 3.5 min for low states and to 1.5 min for high states (Fig. S2) (median values; data from long-block measurements in two animals; non-significant state durations identified by means of permutation and excluded). Of note, responses at the beginning of sensory stimulation blocks were often slightly weaker (initial state low in 78% of all cases).

After tACS, response amplitudes at response-modulated sites remained elevated to levels well above half maximum for periods of up to 10 minutes (Fig. S2) (dominant initial state changed to high (75 % of all cases) before transitioning, within the order of a minute, into a more diffuse low state (Fig. 3E). Accordingly, ratios of mean peak-to-peak response amplitudes for first vs. last 100 trials increased (range 1-2; peak at 1.2-1.4; prior to tACS: range 0.6-1.4; peak at 0.9-1.1) (Fig. 3F), and state-transition frequencies decreased to values below one transition per minute, with a clear peak <0.1 transitions/min covering about 50% of all cases (Fig. 3G).

Still, the ranges of normalized response amplitudes were equal to those at unmodulated sites. This suggests that electrical stimulation had modulated state-dynamics rather than maximum response amplitude (Fig. 3H). Latencies of the audiovisual responses were less variable in the high state (Fig. 3I).

The abovementioned effects of tACS on response amplitude and latency jitter in early vs. late trials, as well as on state-transition frequency were statistically significant (all p≪10^−4^) and large (all |z|>4) (Ranksum test; see Table 2 for details). Taken together, the data consistently point to an increase in local response synchrony within the subpopulation of response-modulated sites, measured across the first 100 trials (but outlasting those, see Figs. 3E and S2) of a respective post-tACS block of audiovisual stimuli. This was further supported by an increase in the power of spike-triggered LFP averages (including spikes from ongoing activity), again indicating an enhancement of local synchrony as an effect of tACS (Fig. S3).

**Table 2.** Pairwise statistics on dynamics of response amplitude time series (control: first-of-day measurement before tACS; post, (un-)mod: post-tACS, (un-)modulated sites; Ranksum tests, Bonferroni-corrected alpha threshold: 0.0167; h=1, significant). Sample sizes: 49 / 142 / 176 control / post, unmod / post, mod.

| condition1 | condition2 | measure | *p* | *h* | *z* | ranksum |
| --- | --- | --- | --- | --- | --- | --- |
| *control* | *post, mod* | means ratio (early vs late): amplitude | $6.9 \times 10^{-23}$ | 1 | -9.8 | 1567 |
| *control* | *post, unmod* |  | 0.1079 | 0 | -1.6 | -1.6 |
| *post, unmod* | *post, mod* | | $<< 10^{-40}$ | 1 | -14.4 | 10895 |
| *control* | *post, mod* | CV-ratio (early vs late): latency | $1.4 \times 10^{-5}$ | 1 | 4.3 | 7289 |
| *control* | *post, unmod* |  | 0.0965 | 0 | -1.7 | 4149 |
| *post, unmod* | *post, mod* | | $3.9 \times 10^{-14}$ | 1 | 7.6 | 28815 |
| *control* | *post, mod* | state-transition frequency | $9.2 \times 10^{-23}$ | 1 | 9.8 | 9449 |
| *control* | *post, unmod* |  | 0.0983 | 0 | 1.7 | 5256 |
| *post, unmod* | *post, mod* | | $3.1 \times 10^{-40}$ | 1 | 13.3 | 33427 |

In the light of these results, we asked whether increases in coherence also occurred in the pre-stimulus activity, indicative of the ongoing network state, at response-modulated sites after tACS offset, and, if so, whether the decline in response amplitude observed after several minutes (Fig. 3E, S2) might be accompanied by a change in coherence as well. To this end, we compared coherence spectra obtained from baseline activity (500ms pre-stimulus window) preceding the first 100, respectively last 100 repetitions of the audiovisual stimulus (Fig. 4). Across the first 100 trials right after tACS, the high response-amplitudes at the response-modulated sites were accompanied by stronger baseline coherence in the theta range (<8Hz) (Fig. 4A), which resembles the abovementioned situation for stimulus-related coherence (Fig. 3D). Coherence at response-modulated sites was also stronger in the beta range around 20Hz, but lower at alpha-band frequencies. During late trials, coherence spectra were rather similar for response-modulated and unmodulated sites (Fig. 4B). Compared to the situation right after tACS, both types of site now showed higher coherence in the alpha range but lower coherence in the gamma-range (Fig. 4C,D). Such a change in gamma-coherence between early and late trials was not observed in the pre-tACS control measurements (evaluated at the sites that were response-modulated after tACS) (Fig. 4E).

**Fig. 4.**
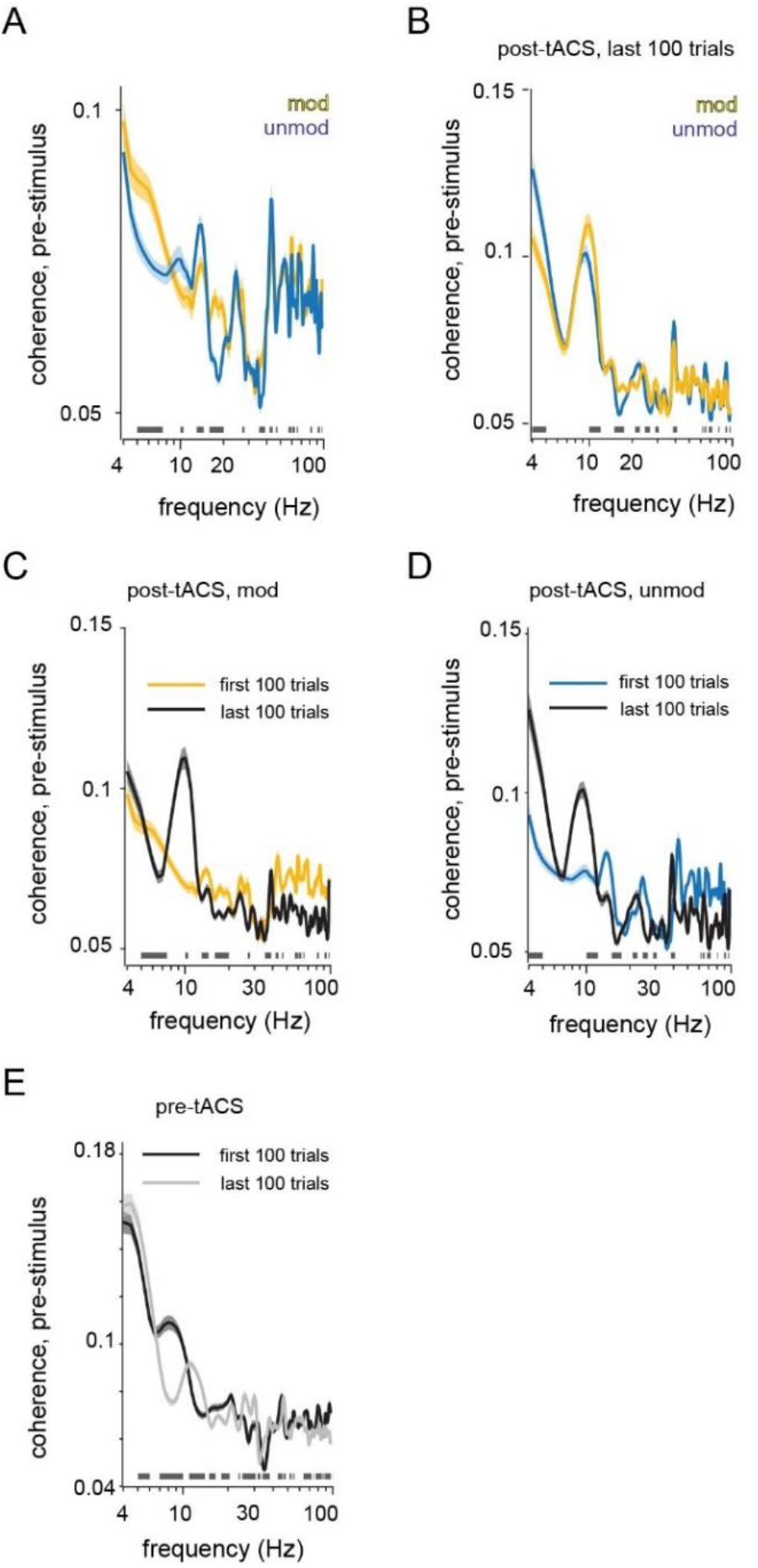
Coherence spectra of pre-stimulus activity for early vs. late trials of audiovisual stimulation. Plots show spectra of mean imaginary coherence between recording sites, quantified for pre-stimulus baseline (500 ms window preceding stimulus onset). **(A)** Coherence during the first 100 trials right after tACS, measured among response-modulated sites (mod) and among unmodulated sites (unmod)., respectively **(B)** Same as (A), but measured between the last 100 trials of the stimulus block. **(C)** Coherence among response-modulated sites during early vs. late trials of the stimulus block after tACS. **(D)** Same as (C), but for unmodulated sites. **(E)** Coherence during early and late trials prior to tACS, same sites as in (E) (data taken from baseline-measurements of the audiovisual responses prior to any tACS in the respective animal). Horizontal bars at frequency axes indicate significance (10.000 permutations). Grey silhouettes indicate SEM.of variation; early, late: first, last 100 trials. Number of recording sites included: pre-tACS control: 49; unmod: 142; mod: 176.

Taken together, the network state right after tACS, when response-modulated sites showed more stable, high response-amplitudes and less variable response latencies, was marked by higher gamma-band coherence, but lower coherence in the theta- / alpha range than during the later state during which responses where more variable also at modulated sites.

Finally, the effect on the time series of response amplitudes and latencies was more often observed after gamma-tACS where it was also covering bigger portions of the cortical surface (Fig. S1) and was longer lasting than when observed following alpha-tACS (Table 1). Effects were observed during times of day spanning widely from 6:00 pm to 3:00 pm, while first-of-day post-tACS measurements were usually performed around 5-6 pm (Fig. S1). Repeated measurements suggest that the number of response-modulated channels substantially decreased from the first to second application of tACS at a respective frequency (Fig. S1), while appearing rather independent from time of day.

## Discussion

We analyzed electrocorticographic recordings from an extended network including occipital, temporal and parietal cortical areas in the ferret, to screen for persistent effects of tACS on intrinsic and stimulus-related activity. Our observations on local synchrony (continuous and spike-triggered power of LFP), synchrony across recording sites (imaginary coherence) and altered dynamics of response-features (peak-to-peak amplitude, extremum latency) consistently point to a synchronizing effect of tACS that can outlast electrical stimulation by at least up to 10 min.

Previous animal studies into the neural underpinnings of tACS were mostly confined to local recordings [17,28] or resorted to regimes that only partly mimic human tACS, in particular more invasive stimulation through electrodes attached to the skull [17] or even implanted into the skull [28], thus bypassing the problem that skin and bone may attenuate current by more than 50% [36]. To our knowledge, no previous work has provided invasive measurements of spatially resolved network activity in a tACS regime that resembles the situation in humans. The present study introduces an approach that fills this gap and yields insights into the neural effects of tACS at the level of large-scale network dynamics.

### Aftereffects of tACS are marked by cross-frequency antagonisms

Notably, we found tACS at gamma-band frequencies to be more effective in persistent modulation of both baseline activity and audiovisual responses compared to alpha-band stimulation. While many previous studies have not considered or not revealed lasting effects of tACS at frequencies other than alpha, e.g. [26], some have provided evidence for antagonistic cross-frequency interactions between tACS-modulated alpha- and gamma-band activity [37,38]. These have been interpreted as enhancement of the intrinsic antagonism between rhythms indicative of functional inhibition [39] (alpha) and functional activation (gamma) [40,41], respectively. In line with the idea of cross-frequency interactions, we observed effects of alpha- and gamma-tACS on prestimulus power and coherence at frequencies other than the targeted (Fig. 2e,f). In particular, gamma-band tACS caused a broadband increase in coherence of baseline activity (indicative of the network state), which was weakest around 10Hz. In turn, stronger gamma-coherence and relatively low alpha-coherence appeared to be related to the state of particularly stable and strong audiovisual responses that we observed predominantly after gamma-tACS (Fig. 4C).

### Plasticity as a candidate mechanism for tACS-aftereffects

The cellular substrates of post-tACS effects remain a subject of intense discussion [13,42]. During current application, high-amplitude tACS that targets endogenous oscillation frequencies was shown to entrain subthreshold potentials and possibly spike patterns [43]. In the present study however, pre-tACS power spectra usually lacked narrow peaks that could have been augmented by the applied tACS frequencies. In addition, sensory stimuli in our approach were separated by randomized intervals (1-2.5 sec), and thus not aligned to the phase of previously applied tACS current, which prevents substantial boost of response amplitudes by phase alignment to entrained endogenous oscillations. Beyond entrainment effects, it has been widely suggested that the coordinated modulation of neuronal excitability may cause plastic changes, e.g. via spike-timing dependent synaptic mechanisms [44,45] which might underlie our present observations as well. By means of an elaborated stimulation protocol, Vossen and colleagues controlled post-tACS increases in alpha power for the possible contribution of entrainment “echoes” [46]. In fact, no entrainment-specific features such as phase-continuity or synchronization to exact stimulation frequencies were observed by Vossen and colleagues, leading to the conclusion that the observed tACS aftereffects on intrinsic activity were most likely plasticity-based. With respect to stimulus processing, a recent EEG study in human subjects [47] has reported significant differences between responses to tactile stimuli recorded prior to and after series of brief intermittent alpha-band tACS (6s, 1mA, several hundred repetitions, frequency tailored to individual’s alpha-peak). From modeling, Sliva and colleagues propose a process of local synaptic facilitation, which may have specifically amplified distal “feedback” input (~70 ms post-stimulus) to the SI circuitry. The difference in response waveform survived averaging over 10 minutes, a time range that matches the maximum duration of effects on response amplitudes in the present study. This again suggests plastic processes as a candidate substrate for the post-tACS effects on stimulus processing we describe here.

## Conclusion

We conclude that prolonged tACS at moderate current intensity and, in particular, at frequencies in the gamma range can exert sustained effects on neural synchrony with characteristics that suggest plasticity-related mechanisms. Subdural, large-coverage recording of neural activity enabled us to identify these effects at the level of an extended cortical network, in a tACS preparation that closely resembles conditions of transcranial electrical stimulation in humans. A deeper understanding of the underlying processes is still required and may benefit from optogenetic or pharmacological interventions to selectively manipulate components of the putative mechanisms. The observed effects on sensory responses promise a means to alter, over extended time periods, aspects of brain function that govern perception and behavior. Follow-up studies, in particular with use of subdural large-scale recordings in chronic preparations, could further promote the development of tACS as a tool to re-adjust brain states in pathological conditions of disturbed functional connectivity [10,12,25].

## Acknowledgment

We thank Dorrit Bystron for her assistance throughout the experiments.

## Abbreviations

tACS: : transcranial alternating current stimulation
pre-tACS: : tags data measured prior to tACS
post-tACS: : tags data measured after tACS
ECoG: : electrocorticography
LFP: : local field potential
mod: : tags data from recording sites at which tACS modulated audiovisual responses
unmod: : tags data from recording sites at which tACS did not modulate audiovisual responses

