## Supplementary material for "Multi-frequency tACS has lasting effects on neural synchrony and audiovisual responses": S1

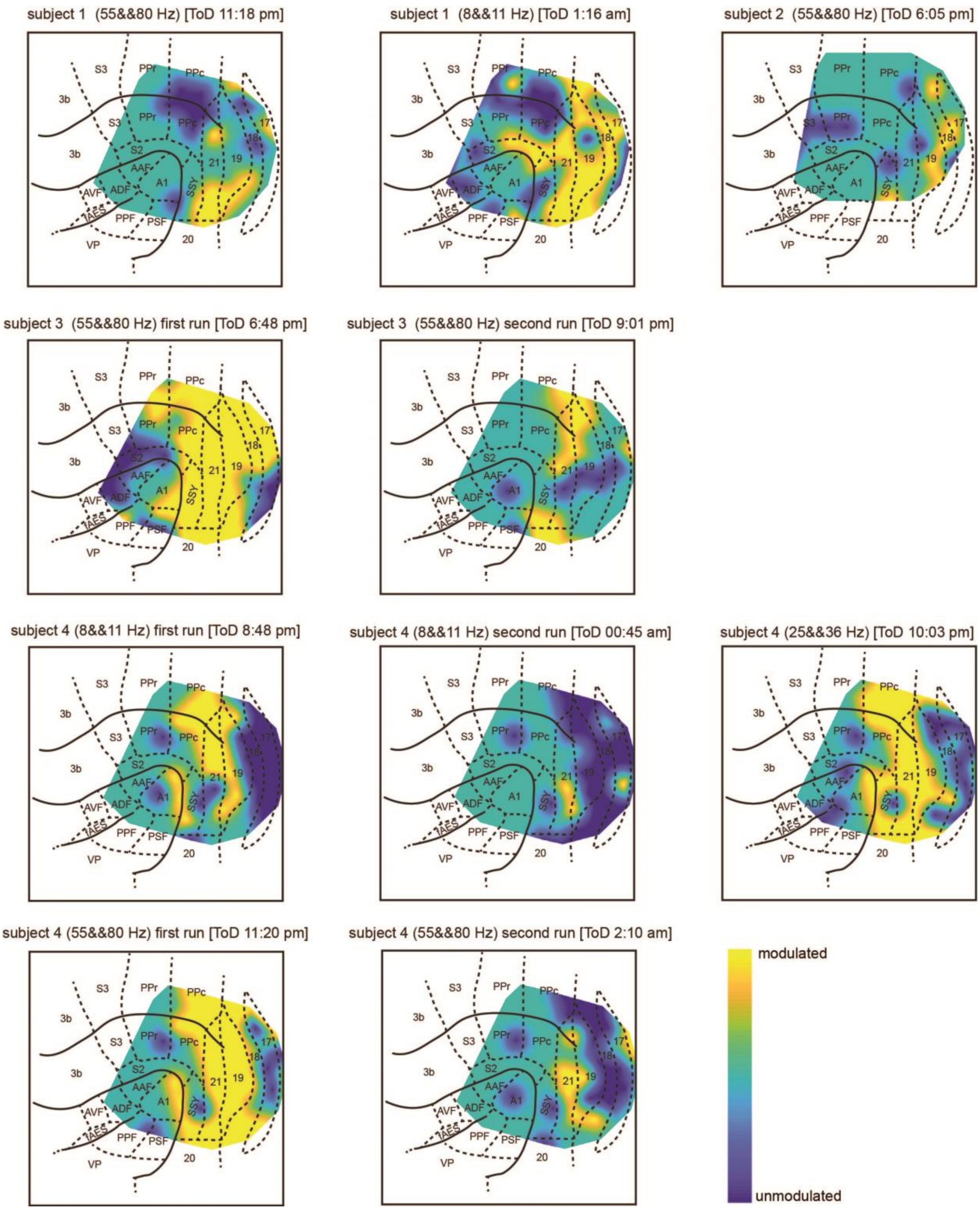


**Fig. S1.** Post-tACS topographies of response-modulated sites. Same logic as for Fig. 3(A-B); frequencies in brackets indicate the two respective sine frequencies from which the tACS current waveform was synthesized. In subjects 3 and 4, where repetitions of the same tACS-band have been run, the first respective application produced substantially more response-modulated sites. Moreover, except for subject 1, gamma-tACS (and possibly beta-tACS, one experiment) appears to produce more response-modulated sites than alpha-tACS.
