## Supplementary material for "Multi-frequency tACS has lasting effects on neural synchrony and audiovisual responses": S2

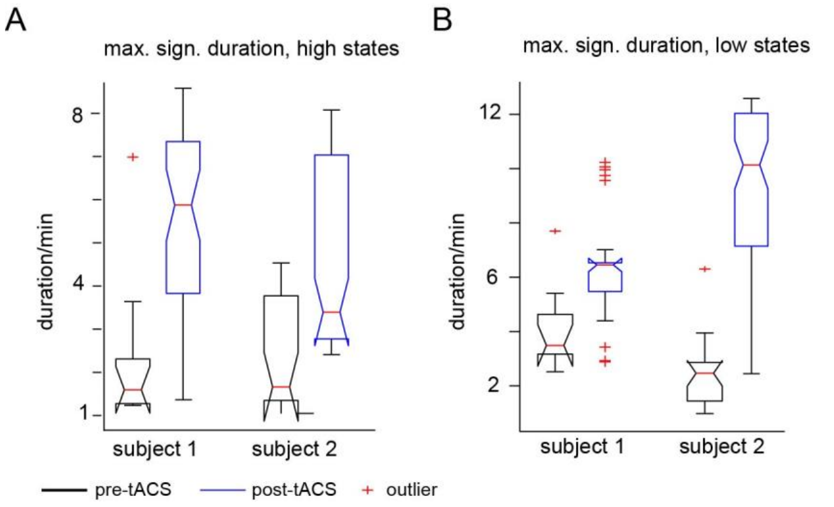


**Fig. S2.** State durations before and after tACS. **(A)** Durations of high-amplitude states as observed in experiments with very long stimulus blocks. Black contour: pre-tACS control measurement; Blue contour: post-tACS measurement, modulated recording sites only. Durations of very short, non-significant fluctuations were excluded (permutation analysis).Note that notches around median indicate 95 % CI. **(B)** same logic for maximum duration of low-amplitude states.
