## Supplementary material for "Multi-frequency tACS has lasting effects on neural synchrony and audiovisual responses": S3

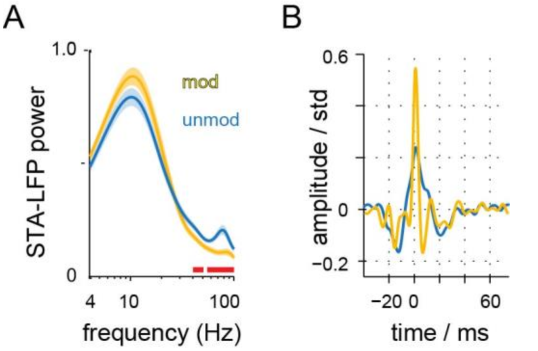


**Fig. S3.** Spike-triggered averages of LFP band activity. **(A)** spike-triggered average of LFP power ( -20ms to +60 ms peri spike); **(B)** grand averages of the spike-triggered LFP waveform.
