## Supplementary material for "Multi-frequency tACS has lasting effects on neural synchrony and audiovisual responses": S4

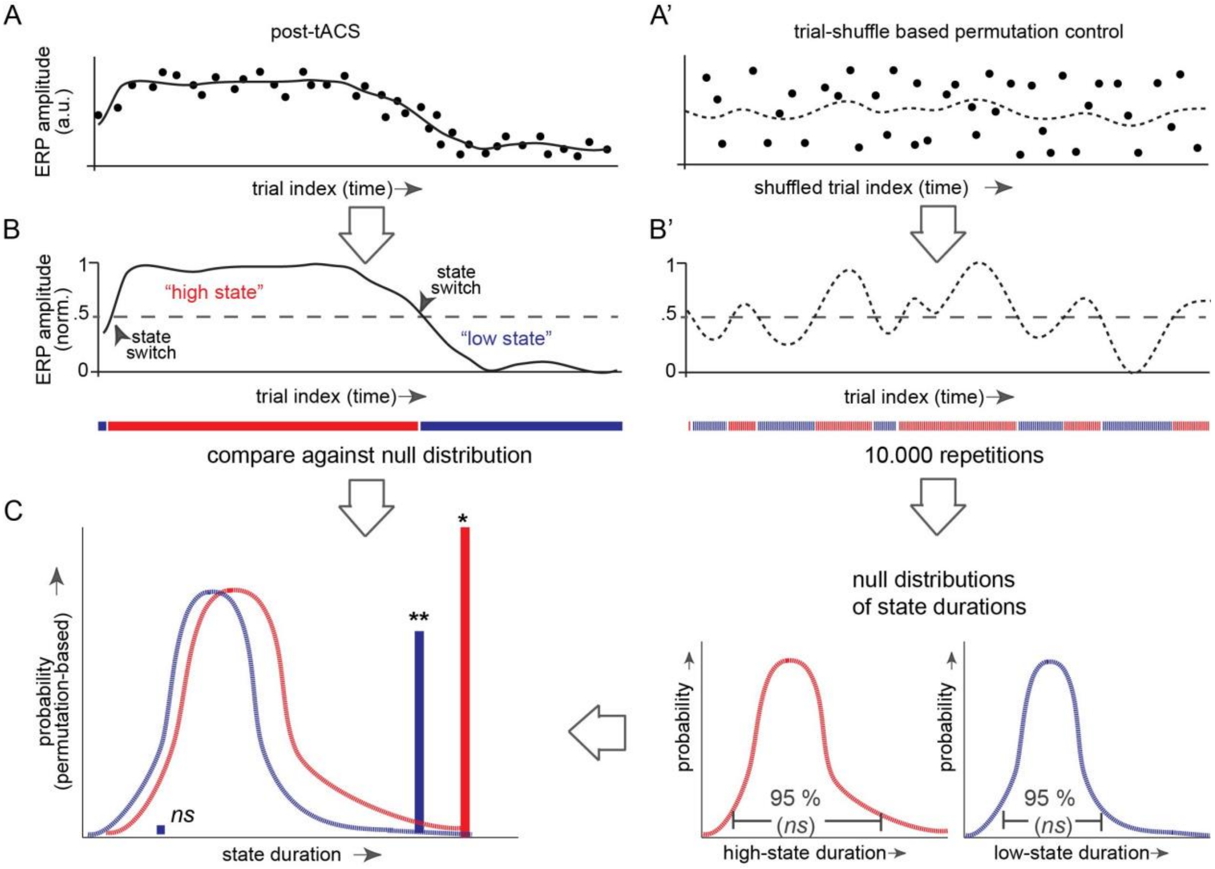


**Fig. S4.** Methods illustration: analysis of state dynamics in ERP amplitude-time series. **(A)** Illustration of a typical time series of ERP amplitudes (filled circles) as observed at a tACS modulated recording site. To further evaluate state dynamics, the series is first smoothed by convolution with a 25-elements boxcar kernel. **(B)** to assign each trial's ERP amplitude to either "high state" or "low state", the amplitude range of the convolved series is re-scaled to [0,1] and values above (below) a normalized amplitude of 0.5 are then classified as "high state" ("low state"). Periods of continuous high (low) states are indexed by red (blue) horizontal bars below. **c** to test the duration of each such period for statistical significance, durations are finally compared to permutation-based null distributions that are obtained based on trial-shuffling the empirical data **(A',B')**.
